# Multi-step regulation of transcription kinetics explains the non-linear relation between RNA polymerase II density and mRNA expression in Dosage Compensation

**DOI:** 10.1101/203596

**Authors:** Pouria Dasmeh

**Affiliations:** Departement de Biochimie, Université de Montréal, 2900 Edouard-Montpetit, Montreal, Quebec H3T 1J4, Canada.; Centre Robert Cedergren en Bioinformatique et Génomique, Université de Montréal, 2900 Edouard-Montpetit, Montreal, Quebec H3T 1J4, Canada.; Department of Chemistry and Chemical Biology, Harvard University, Cambridge, MA USA 02139.

## Abstract

In heterogametic organisms, expression of unequal number of X chromosomes in males and females is balanced by a process called dosage compensation. In *Drosophila* and mammals, dosage compensation involves nearly two-fold up-regulation of the X chromosome mediated by dosage compensation complex (DCC). Experimental studies on the role of DCC on RNA polymerase II (Pol II) transcription in mammals disclosed a non-linear relationship between Pol II densities at different transcription steps and mRNA expression. An ~20-30% increase in Pol II densities corresponds to a rough 200% increase in mRNA expression and two-fold up-regulation. Here, using a simple kinetic model of Pol II transcription calibrated by *in vivo* measured rate constants of different transcription steps in mammalian cells, we demonstrate how this non-linearity can be explained by multi-step transcriptional regulation. Moreover, we show how multi-step enhancement of Pol II transcription can increase mRNA production while leaving Pol II densities unaffected. Our theoretical analysis not only recapitulates experimentally observed Pol II densities upon two-fold up-regulation but also points to a limitation of inferences based on Pol II profiles from chromatin immunoprecipitation sequencing (ChIP-seq) or global run-on assays.

---

Unequal number of X chromosomes in males and females of several organisms imposes a dosage problem on expression of X-linked genes. In the absence of a proper regulatory mechanism, this imparity potentially leads to unequal expression of X-linked genes and sex lethality. To overcome this challenge, a “dosage compensation” mechanism is evolved to compensate the expression of X chromosomes^1–3^. In Drosophila, the one copy of X-chromosome in males is roughly transcribed by twofold to balance the expression of the two X chromosomes in females^4–7^. In mammals, an X chromosome in females is primarily inactivated to balance the expression of X-linked genes with males^8–10^. Moreover, mammalian X chromosome is further hypertranscribed in order to satisfy X-Autosome expression ratio^11,12^.

Recent studies in Drosophila and mammals using chromatin immunoprecipitation and global-run-on sequencing (i.e., ChIP-seq and GRO-seq) have addressed the interference of dosage compensation with Pol II transcription at different steps. In mouse embryonic stem cells (ES cells), dosage compensation is shown to increase Pol II densities at initiation (i.e., phosophorylated Pol II at Serine 5, i.e., Pol II-S5P) without significant changes in the elongated form of Pol II (i.e., phosophorylated Pol II at Serine 2 or Pol II-S2P)^13^. In another study of dosage compensation in mouse female embryonic kidney fibroblasts, both Pol II-S5P and PolII-S2P densities were found enhanced^12^. In Drosophila, whether dosage compensation facilitates Pol II progression across active X-linked genes^14^ or enhance recruitment of Pol II to promoters^15^ has been controversial^16–18^.

From the above-mentioned experimental studies, one emerging pattern is a nonlinear relationship between Pol II densities at different steps of transcription and mRNA expression level**Error! Bookmark not defined.**. In Drosophila, Pol II tag density over the bodies of X-linked genes compared to autosomal genes is shown to differ by a factor of ~1.4^14^ with ~1.2 folds increase at promoters^17^. In the case of X chromosome up-regulation in mammals, Pol II at promoters and along the body of active genes was reported to be increased by ~1.3 and ~1.2 fold respectively^12^. In both examples, mRNA levels are increased by ~two-fold upon hyper transcription. How does an ~30% increase in Pol II density gives rises to ~200% increase in mRNA production?

Here, we justify this non-linear relationship based on multi-step regulation of transcription machinery. Our assumption is that dosage compensation is achieved by proper alterations of different steps of Pol II transcription. We use the following kinetic framework for Pol II transcription (Figure 1A):

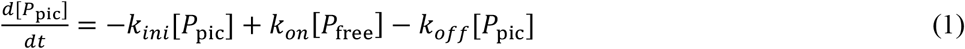

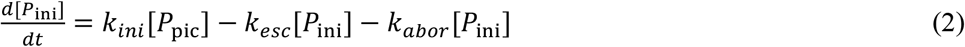

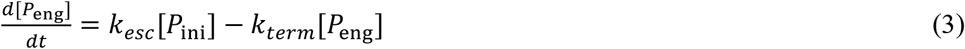

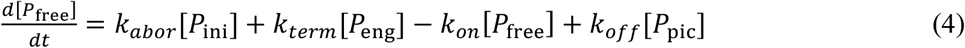

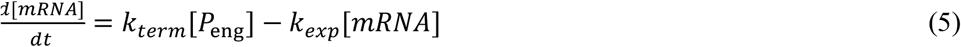

**Figure 1.**
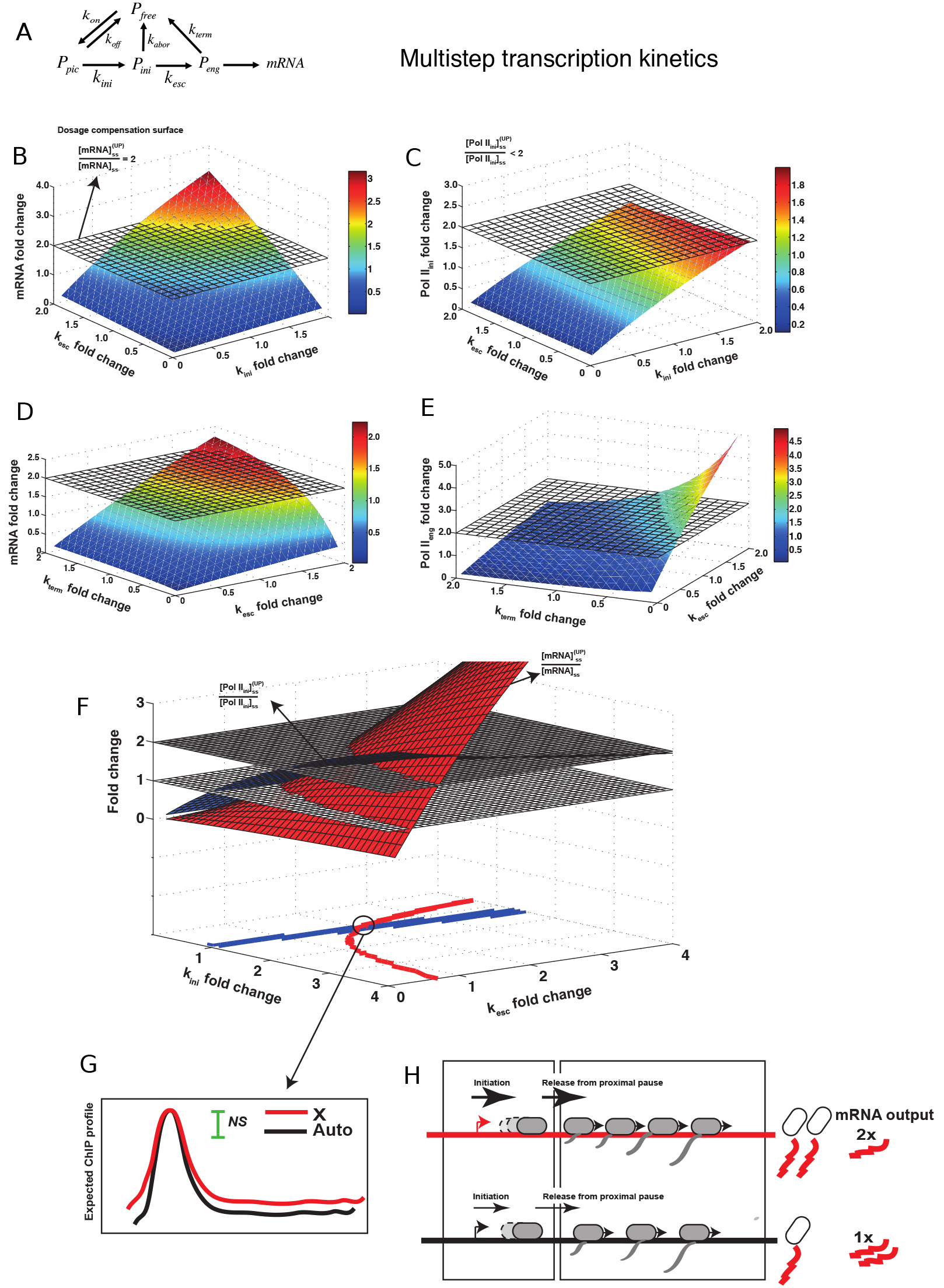
A) RNA polymerase II transcription can be perturbed at different steps (equations 1-6) to give two-fold mRNA production. In B) 3D curve for mRNA fold change is represented as a function of initiation, *k*_ini_, and promoter-escape, *k*_esc_, rate constants. C) 3D curve for *P*_ini_ fold changes as a function of k_ini_ and k_esc_. D) mRNA fold change as a function of *k*_esc_ and elongation-termination rate constant, k_term_. E) 3D curve for *P*_eng_ change as a function of k_esc_ and k_term_. F) 3D curves for fold changes in mRNA (red) and P_ini_ (blue) are crossed with fold change=2 and fold change=1 planes respectively with their projection onto a 2D *k*_ini_-*k*_esc_ plane shown in red and blue. G) ChIP profile for Pol II transcription can be misinterpreted as no significant change in *P*_ini_ and significant changes in *P*_eng_ while, as shown schematically in H) both *k*_ini_ and *k*_esc_ are increased at the same time.

Equations 1-5 describe dynamics of Pol II at different transcriptional steps (i.e, in pre-initiation complex, *P*_pic_, at initiation, *P*_ini_, engaged to gene bodies, *P*_eng_, and as free molecules, *P*_free_) and mRNA molecules. In this framework, free Pol II molecules bind to and unbound from promoters with the rate constants *k*_on_ and *k*_off_ and proceed to initiation and elongation with the rate constants of *k*_ini_ and *k*_esc_. In addition to termination described by the rate constant *k*_term_, Pol II transcription is stopped by abortive initiation with the rate constant *k*_abor_. We modeled mRNA production and clearance by the term (*k*_term_[*P*_eng_] − *k*_exp_[*mRNA*]) simplifying splicing and mRNA export out of nucleus into one rate constant, *k*_exp_.

At steady-states, the relation between steady state mRNA expression and Pol II abundance at different steps of transcription can be expressed as:

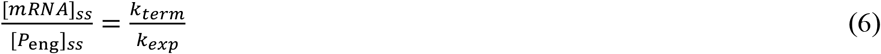

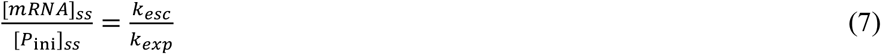

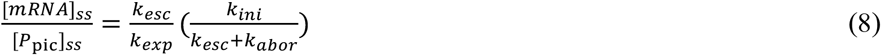

The ratios of Pol II abundances at different steps upon up-regulation (denoted by an “Up” superscript) to the original system are written as:

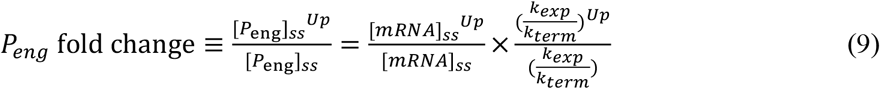

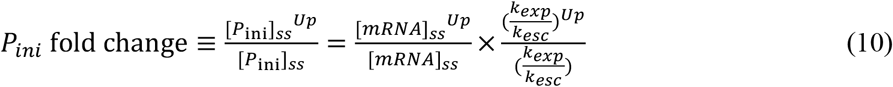

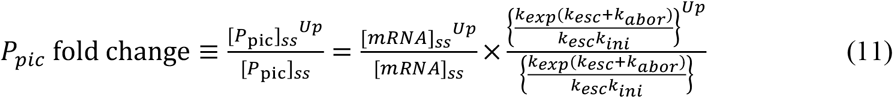

From equations 9 to 11, any change in Pol II abundance is proportional to [mRNA] fold change (i.e.,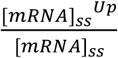) scaled with the ratio of up-regulated rate constants to the original ones (e.g., 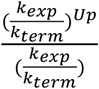 in the case of *P*_eng_ fold change). To calculate the left sides of equations 9-11 and to check whether we can reckon the experimentally observed increased Pol II density at promoters and along the gene bodies, we calculated values of new rate constants assuming two-fold up-regulation. Theoretically, up-regulation can be modeled by one-step (i.e., changing one rate constant), two-step or multi-step perturbation to the original transcription system defined by equations 1-5. Once dosage compensation was achieved (i.e., 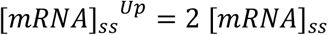), the new rate constants were used to calculate the fold changes in Pol II density at different stages of transcription using equations 9 to 11. We employed *in vivo* estimates of Pol II transcription rate constants in mammalian cells (See Table S1 for parameters used in the model)^19^ and for simplicity assumed no change in mRNA export rate upon dosage compensation (i.e., 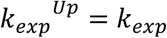).

Perturbing one transcription step to achieve two-fold mRNA production linearly increases Pol II abundance in subsequent steps (See Table S2). For example, increasing the rate of initiation (i.e., by increasing k_ini_) results in increased densities of *P*_ini_, and *P*_eng_. However, two-step perturbation of transcription machinery causes less than two-folds increase in abundance of Pol II at any step which its production and clearance rate are increased simultaneously (See Table S3 and Figures S1-S10 for details). Figure 1B shows mRNA fold change as a function of *k*_ini_ and *k*_esc_. The intersection of the plane 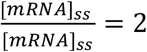 with 3D curve of mRNA fold change defines a dosage compensation surface where combinations of *k*_ini_ and *k*_esc_ result in two-fold hyper transcription. From the figure, mRNA production is doubled by increasing initiation and promoter-escape rate by ~1.2 to 2 folds, simultaneously. However, as shown in Figure 1C, *P*_ini_ is clearly enriched less than two fold when *k*_ini_ and *k*_esc_ are changed at the same time. Given the original kinetic rate constants, ~30% increase in *P*_ini_ corresponds to two-fold mRNA production. A similar situation holds for P_eng_ when k_esc_ and k_term_ are perturbed at the same time (Figures 1D and 1E). An ~20% increase in Pol II abundance along the gene bodies is associated with two-fold mRNA production.

Although perturbing initiation and promoter-escape rates gives rises to *P*_ini_ fold changes in agreement with experimentally observed values, *P*_eng_ is increased by~two-fold (See Table S3 for Pol II abundance at gene bodies while k_ini_ and k_esc_ are perturbed). We thus checked a three-step perturbation analysis and found a combination of k_ini_, k_esc_ and k_term_ that satisfies ~10% to 30% increase in *P*_ini_ and gene bodies upon dosage compensation (See Table S4 and S5 for details).

Next, we asked whether multi-step regulation of transcription can account for two-fold mRNA expression while *P*_ini_ is unaffected. In Figure 1F, red and blue 3D curves show mRNA and *P*_ini_ fold changes as functions of *k*_ini_ and *k*_esc_. The intersection of mRNA 3D curve with the plane at fold change=2 defines a 2D curve for dosage compensation which is projected on *k*_ini_;-*k*_esc_ plane in Figure 1F. We also projected the intersection of 3D *P*_ini_ levels with the plane at fold change=1 in blue. These two curves crossed each other at 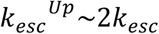 and 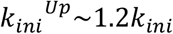 causing two-fold mRNA production and insignificant changes in *P*_ini_ levels. This condition corresponds to an expected ChIP profile in Figure 1G which is most likely misinterpreted as no change in Pol II densities at initiation and a significant change in Pol II densities at elongation steps, although both steps have been enhanced (schema in Figure 1H). In line with previous studies on erroneous inferences from ChIP profiles^20^ and inapplicability of ChIP-seq and Gro-seq in study of Pol II turnover^21^, our study systematically shows the limitation of these methods in addressing relevance of Pol II enrichment at different transcription steps.

Taken these together, our theoretical approach suggests that Pol II transcription is most likely regulated at multiple steps in dosage compensation. How is a multi-step regulation modulated by DCC? There is compelling evidence that DCC proteins, individually or in synergy, influence different transcription steps^22^. For example, in mammals it is shown that MSL1 and MOF, two members of DCC complex, contribute to enhanced densities of Pol II-S5P and therefore facilitates initiation^13^. In addition, MOF as an acetyltransferase is responsible for H4K16ac, a histon modification which decompacts nucleosomes and enhanced promoter-escape and transcriptional elongation^14,18^. As we showed in this work, enhancing initiation, promoter-escape and elongation rates suffice to explain the nonlinearity between Pol II levels and mRNA expression.

The changes in kinetic constants are essential in reproducing the patterns of Pol II transcriptional regulation as shown in this work. Two approaches can be used to measure and compare kinetic rate constants. First, fluorescence recovery after photobleaching (FRAP) experiments^19^ can be used to infer kinetic rate constants of Pol II transcription in selected X-linked and autosomal genes in Drosophila or in mammalian cells. Second, following the work by Kim and Marioni^23^, kinetic rate constants can be inferred from RNA-seq data of Pol II transcription at individual steps. We anticipate future experiments to address these issues.

## Acknowledgement

Author is grateful to professor Asifa Akhtar and Max Planck Society for providing a research fellowship in Max Planck Institute for Immunobiology and Epigenetics, Freiburg, Germany.

## Supplementary Information

**Table S1.**
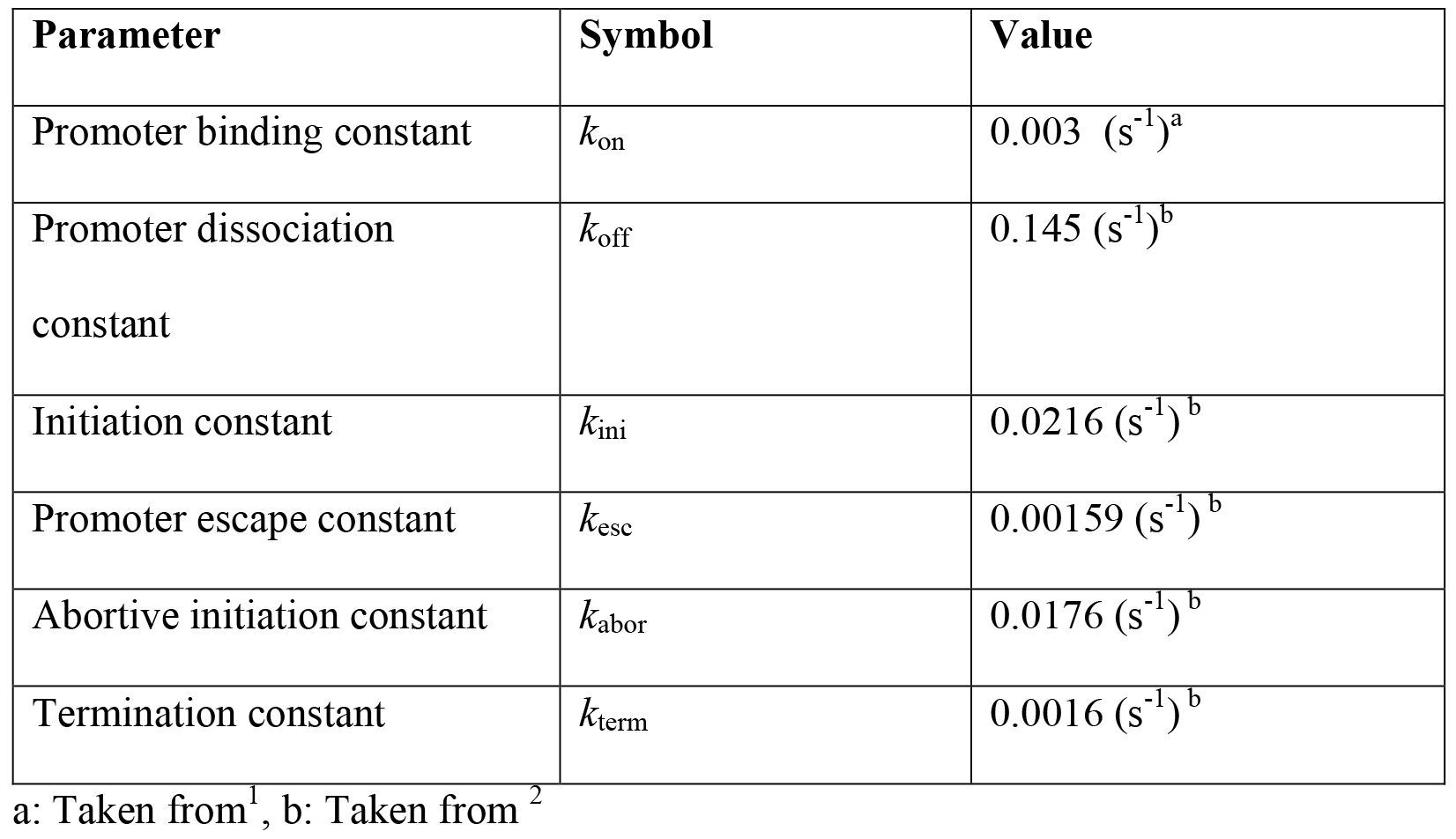
Kinetic constants used in the kinetic model for POL-II transcription.

| Parameter | Symbol | Value |
| --- | --- | --- |
| Promoter binding constant | $k_{\text{on}}$ | $0.003 \text{ (s}^{-1}\text{)}^{\text{a}}$ |
| Promoter dissociation constant | $k_{\text{off}}$ | $0.145 \text{ (s}^{-1}\text{)}^{\text{b}}$ |
| Initiation constant | $k_{\text{ini}}$ | $0.0216 \text{ (s}^{-1}\text{)}^{\text{b}}$ |
| Promoter escape constant | $k_{\text{esc}}$ | $0.00159 \text{ (s}^{-1}\text{)}^{\text{b}}$ |
| Abortive initiation constant | $k_{\text{abor}}$ | $0.0176 \text{ (s}^{-1}\text{)}^{\text{b}}$ |
| Termination constant | $k_{\text{term}}$ | $0.0016 \text{ (s}^{-1}\text{)}^{\text{b}}$ |

a: Taken from<sup>1</sup>, b: Taken from<sup>2</sup>

**Table S2a.**
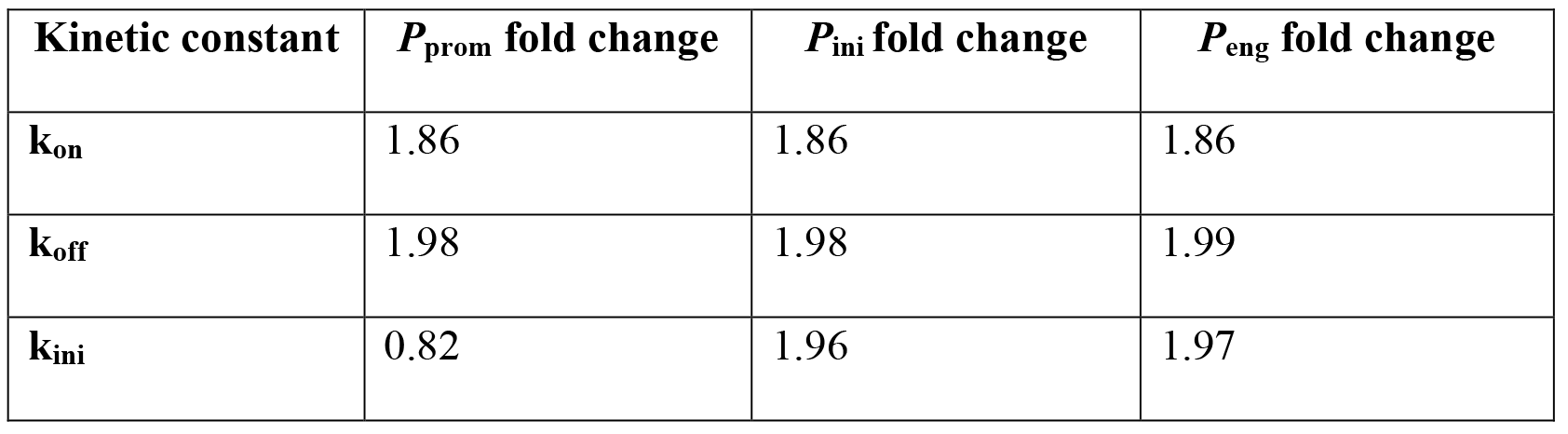
The fold change of POL-II at promoters. initiation and engaged phase in dosage compensated genes (1.8 < *mRNA* fold change < 2.2) caused by relative changes in each kinetic constant

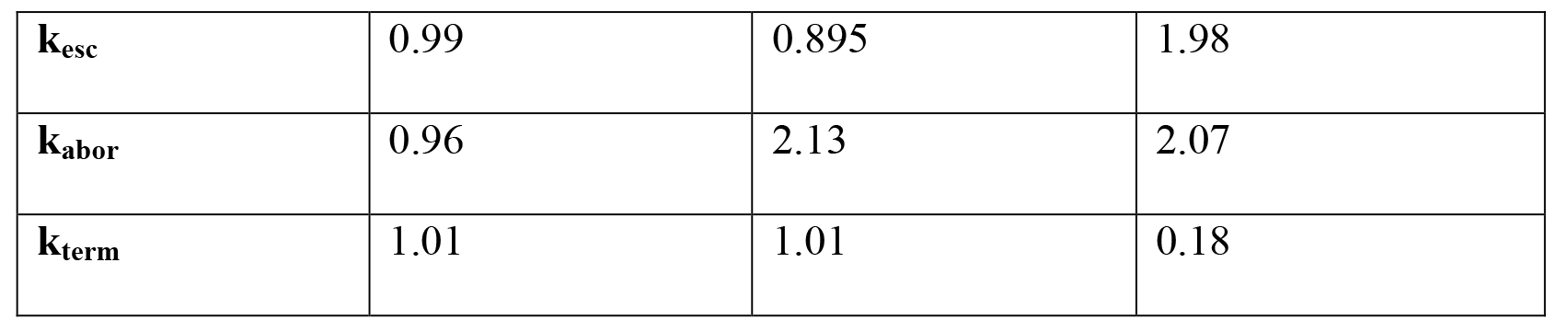

**Table S3.**
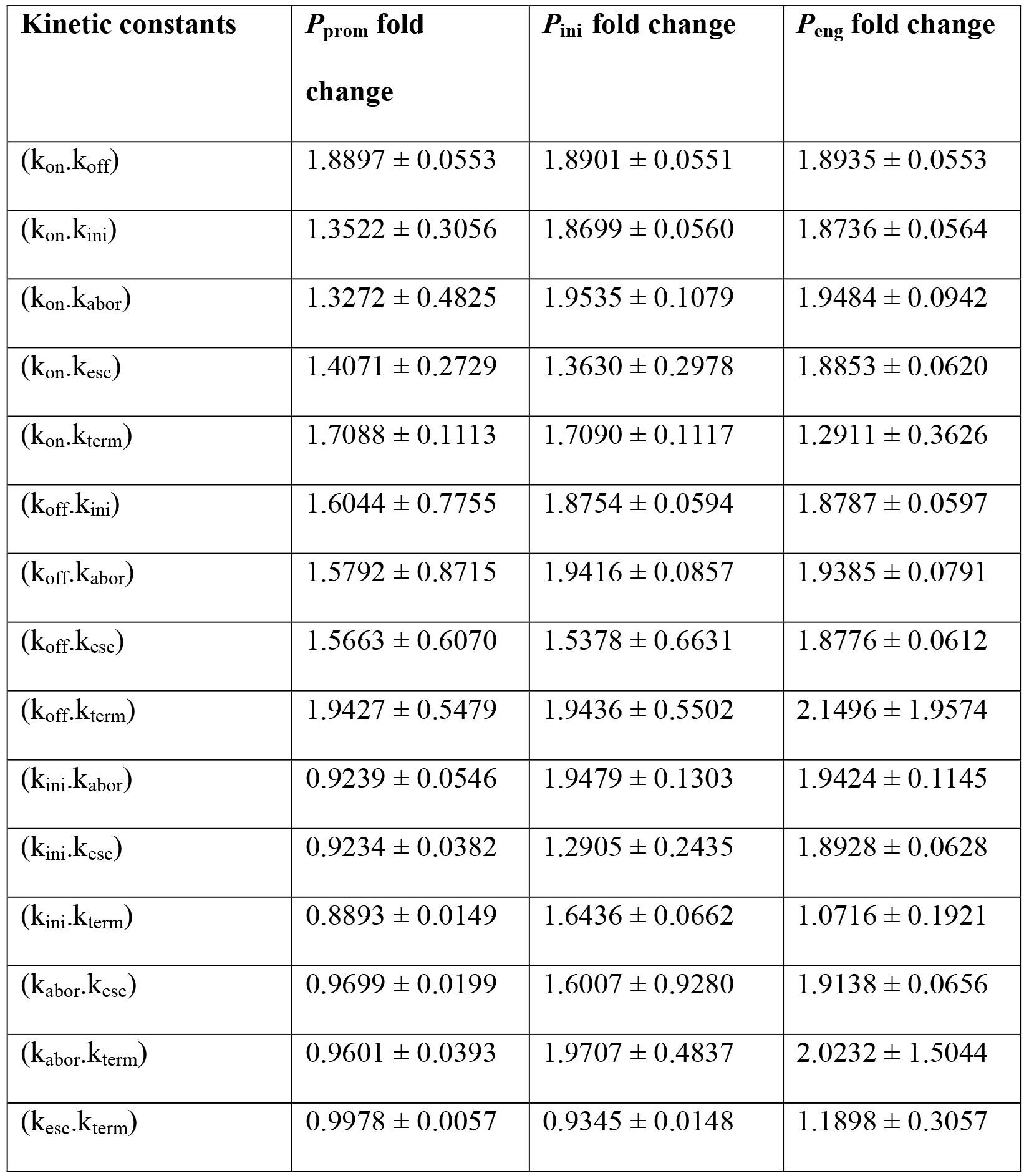
Results of the fold change of POL-II at promoters. initiation and engaged phase in dosage compensated genes (X-chromosome) when 1.8 < *mRNA* fold change < 2.2.

**Table S4a.**
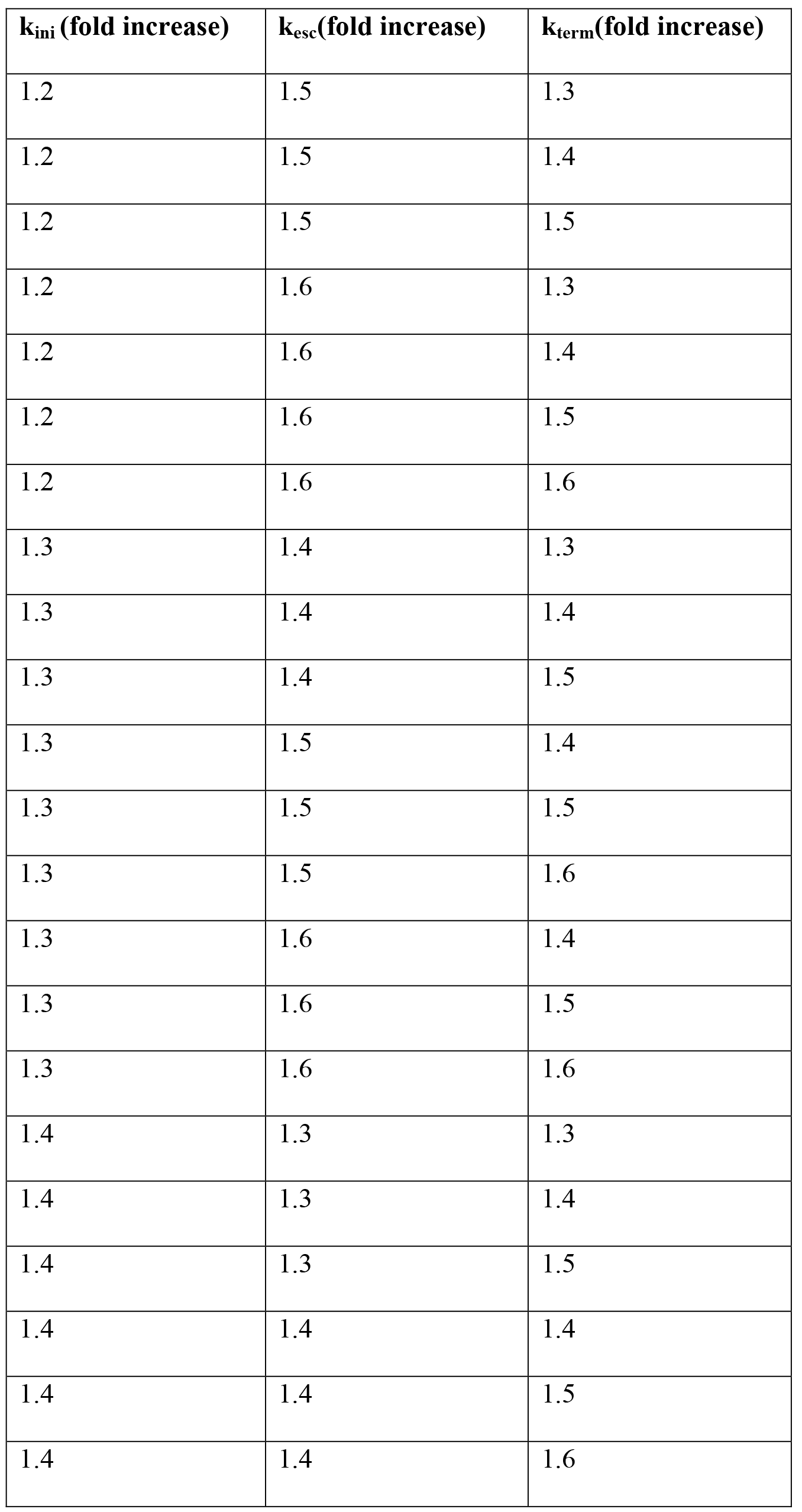
Different combinations of rate constants which satisfy 1.1<*P*_eng_ fold change<1.3. 1.1<*P*_ini_ fold change<1.3 and 1.1<*P*_prom_ fold change<1.3.

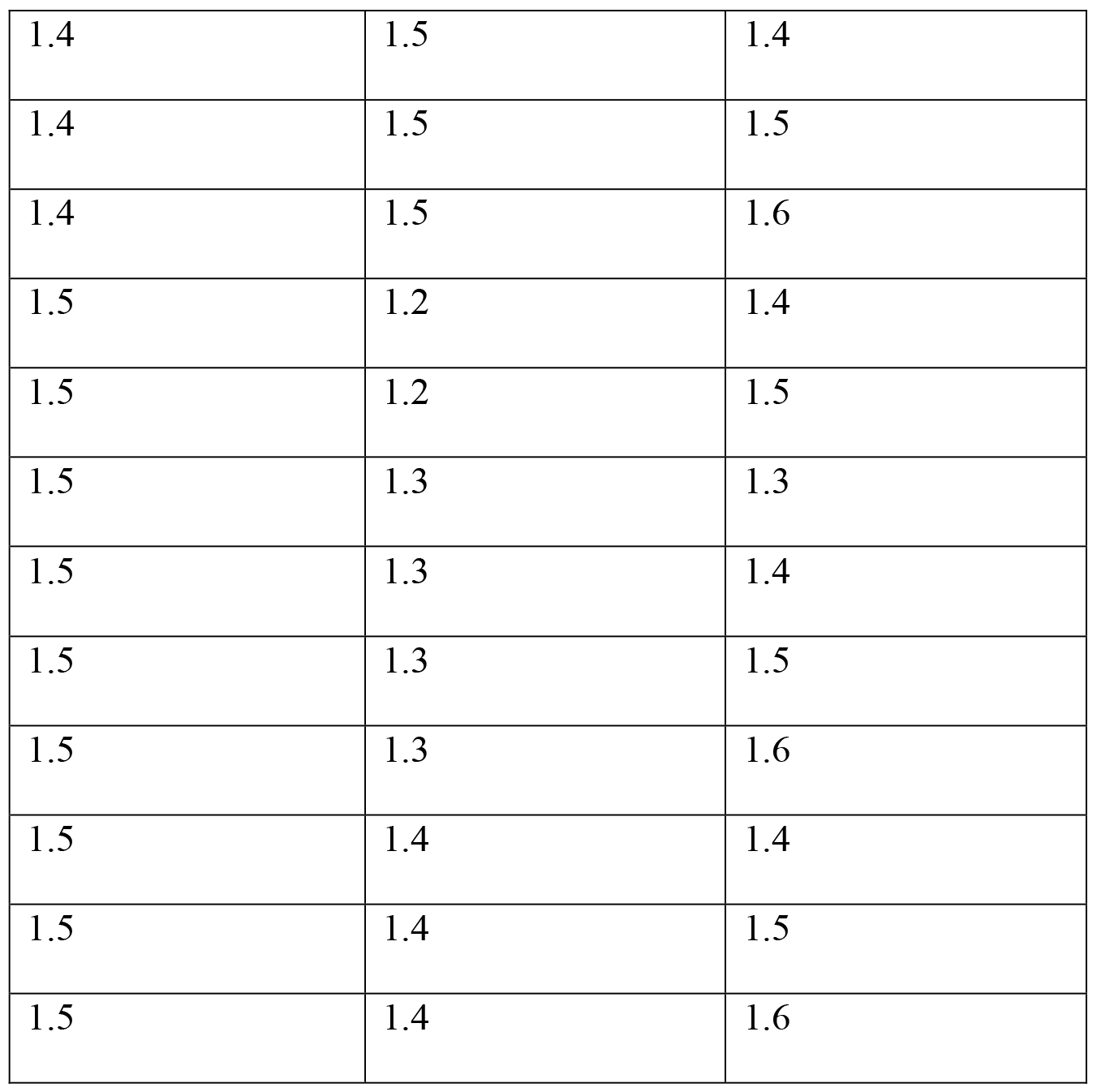

The correlations between the kinetic constants and POL-II abundance at different stages of transcription are shown in Table S5.

**Table S5.** The correlations (+) or anticorrelation (−) between rate constants and POL-II abundance at different stages of transcription.

| **Parameter** | **$P_{\text{prom}}$** | **$P_{\text{ini}}$** | **$P_{\text{prom}}+P_{\text{ini}}$** | **$P_{\text{eng}}$** |
| --- | --- | --- | --- | --- |
| **$k_{\text{on}}$** | + | + | + | + |
| **$k_{\text{off}}$** | - | - | - | - |
| **$k_{\text{ini}}$** | - | + | + | + |
| **$k_{\text{abor}}$** | + | - | - | - |
| **$k_{\text{esc}}$** | - | - | - | + |
| **$k_{\text{term}}$** | + | + | + | - |

In the following figures, the effect of increase or decrease in rate constants on mRNA production is quantified in terms of *mRNA ratio* defined as:

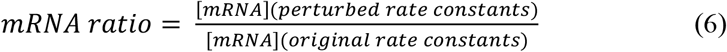

Other abundance ratios defined in this work (e.g., *P*_prom_ *ratio*) are calculated similarly (i.e., 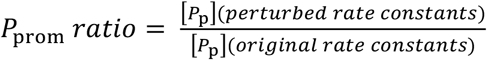).

**Figure S1.**
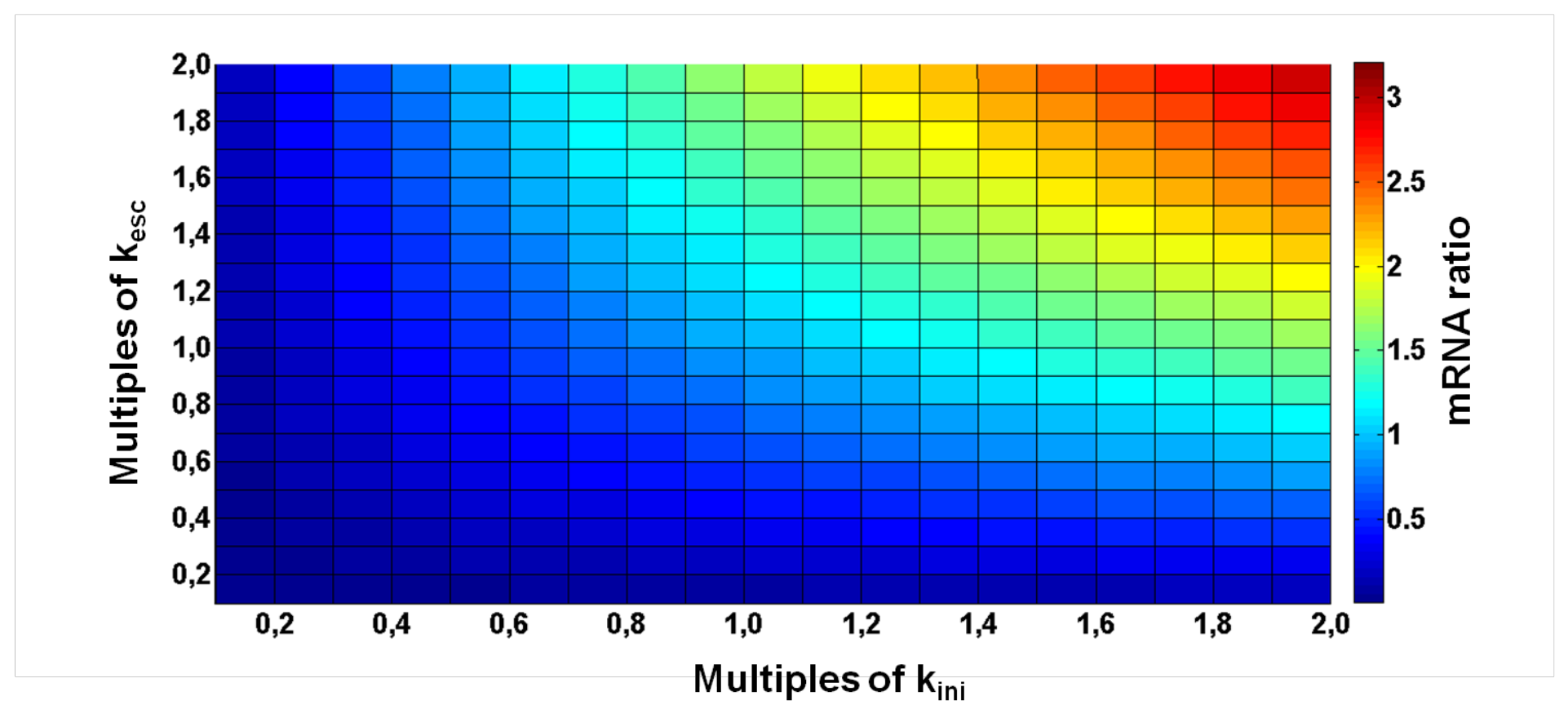
*mRNA* fold change as a function of varying k_ini_ and k_esc_ from one-tenth to two-fold of their original values (i.e., k_ini_=0.0216 s^−1^ and kesc=0.00159 s^−1^).

**Figure S2.**
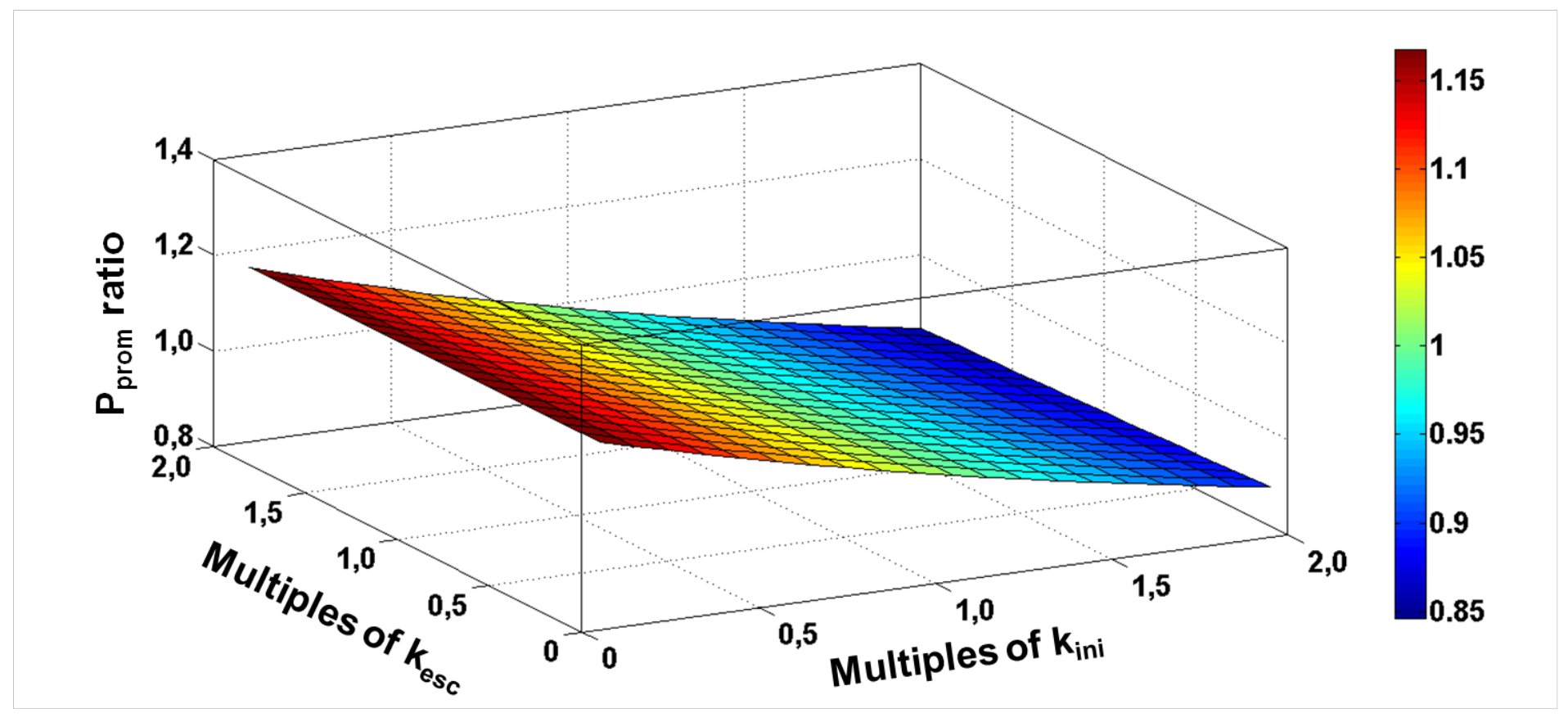
*P*_prom_ fold change as a function of varying k_ini_ and k_esc_ from one-tenth to two-fold of their original values (i.e., k_ini_=0.0216 s^−1^ and kesc=0.00159 s^−1^).

**Figure S3.**
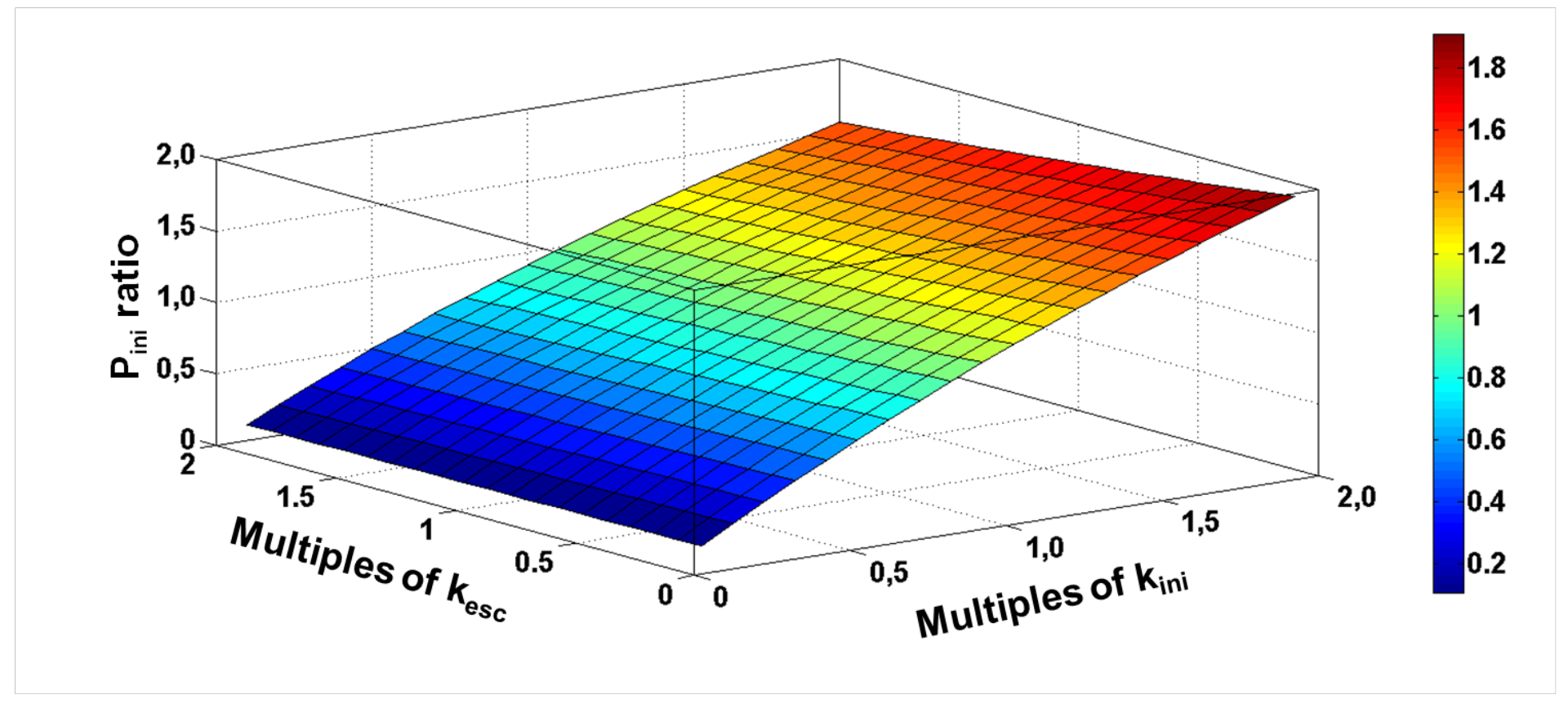
*P*_ini_ fold change as a function of varying k_ini_ and k_esc_ from one-tenth to two-fold of their original values (i.e., k_ini_=0.0216 s^−1^ and kesc=0.00159 s^−1^).

**Figure S4.**
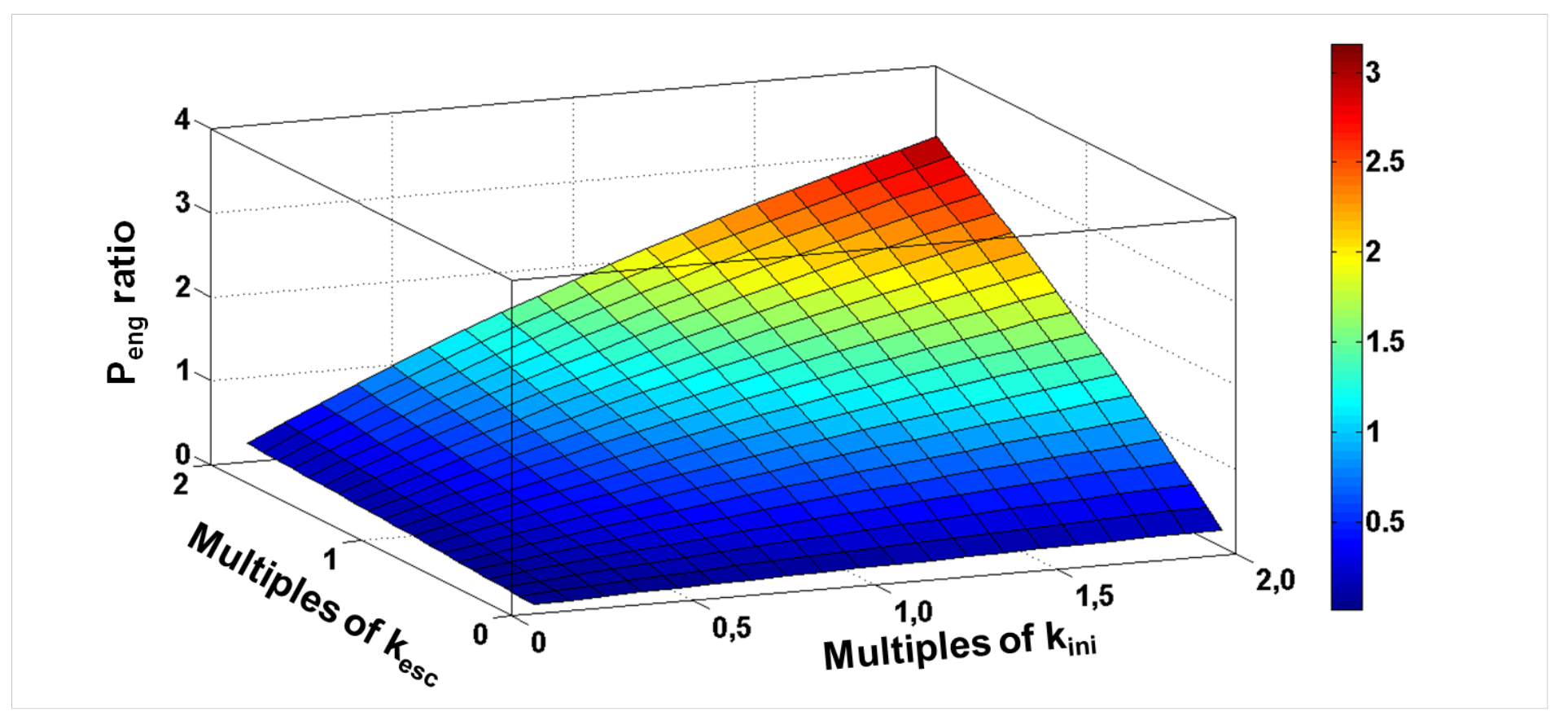
*P*_eng_ fold change as a function of varying k_ini_ and k_esc_ from one-tenth to two-fold of their original values (i.e., k_ini_=0.0216 s^−1^ and kesc=0.00159 s^−1^).

**Figure S5.**
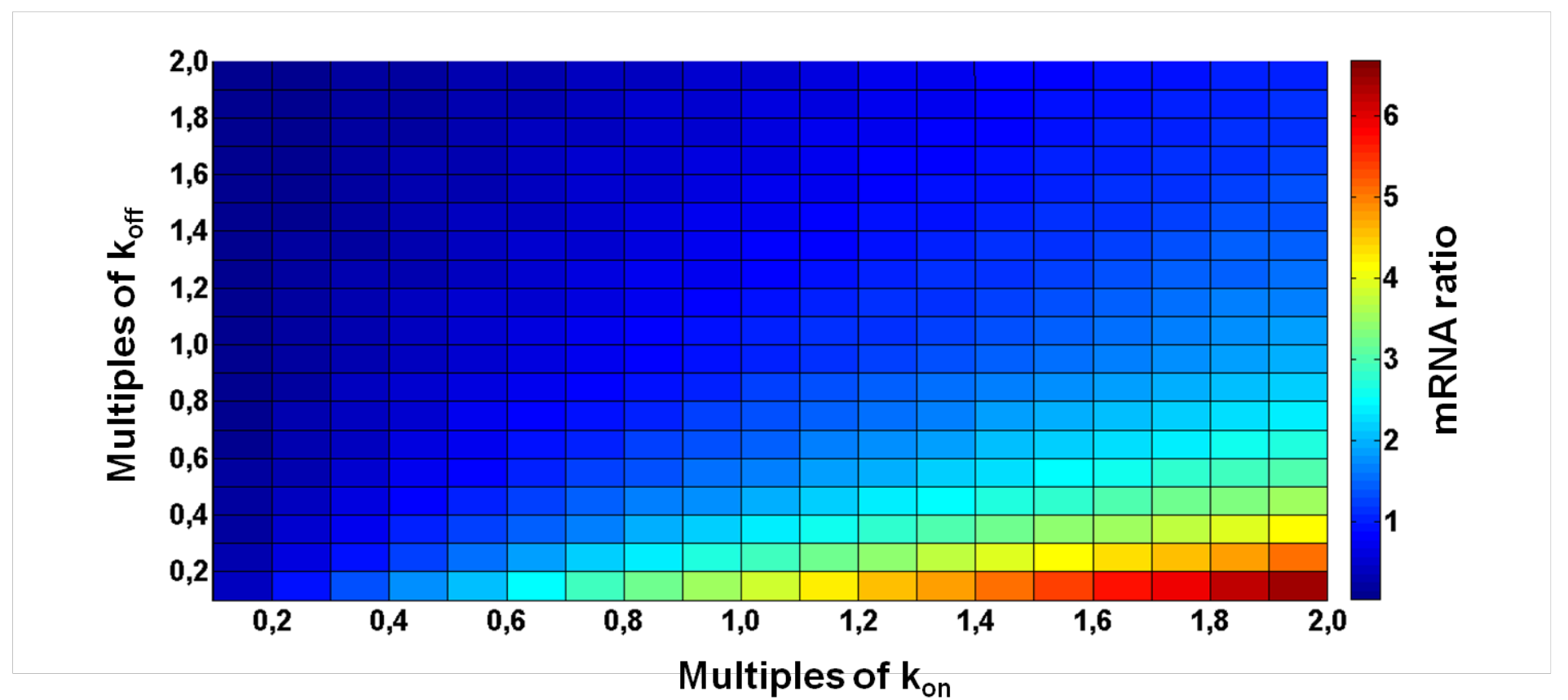
*mRNA* fold change as a function of varying k_on_ and k_off_ from one-tenth to two-fold of their original values (i.e., k_on_=0.0216 s^−1^ and k_off_=0.00159 s^−1^).

**Figure S5.**
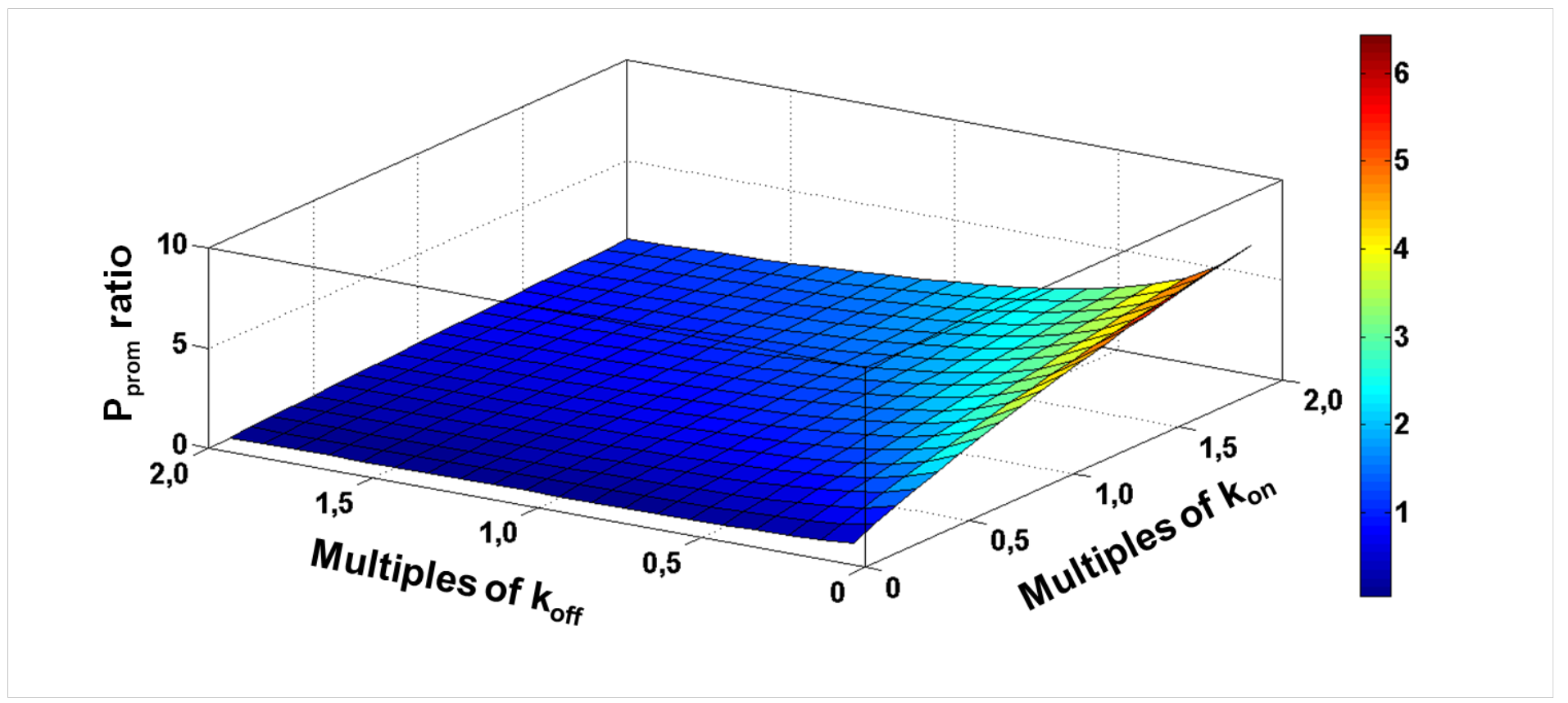
*P*_prom_ fold change as a function of varying k_on_ and k_off_ from one-tenth to two-fold of their original values (i.e., k_on_=0.0216 s^−1^ and k_off_−0.00159 s^−1^).

**Figure S6.**
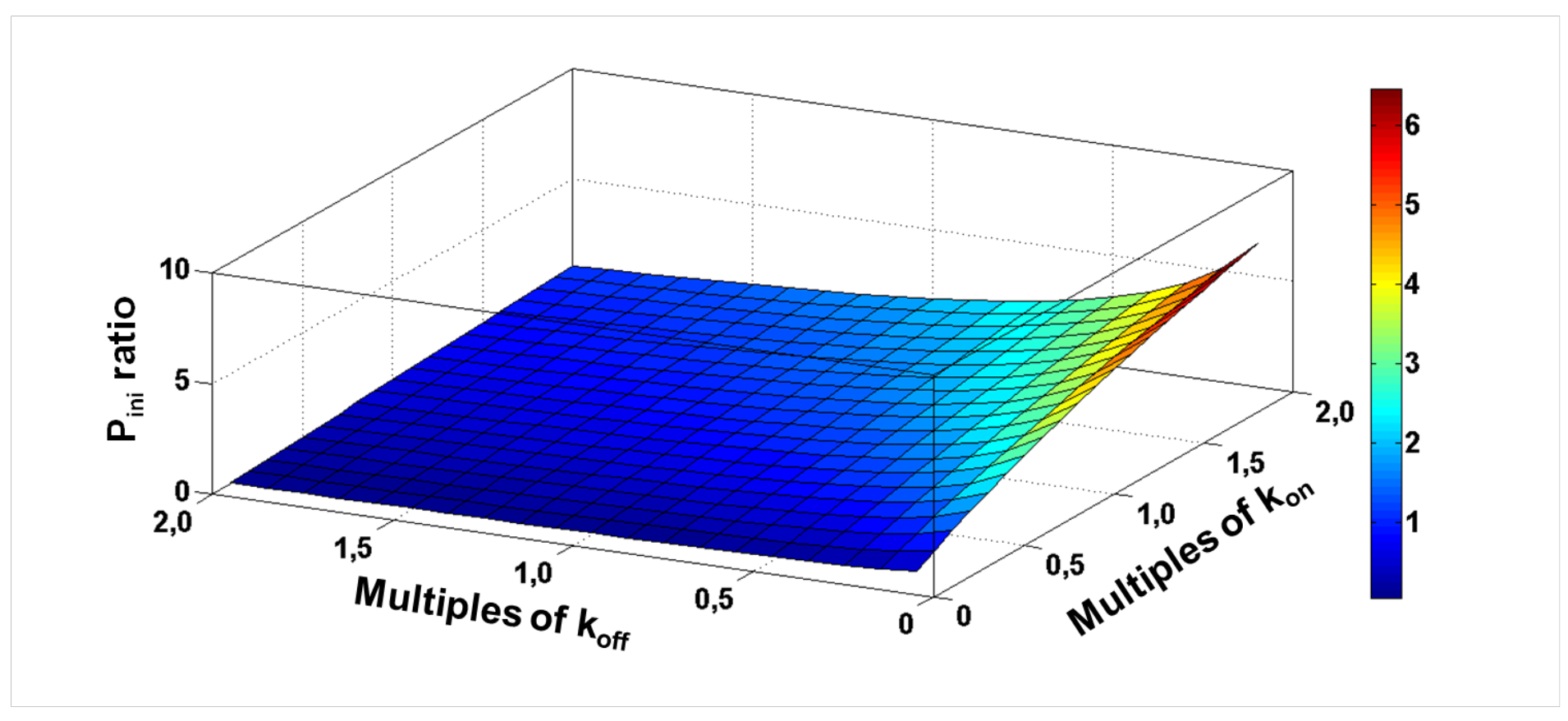
*P*_ini_ fold change as a function of varying k_on_ and k_off_ from one-tenth to two-fold of their original values (i.e., k_on_=0.0216 s^−1^ and k_off_=0.00159 s^−1^).

**Figure S6.**
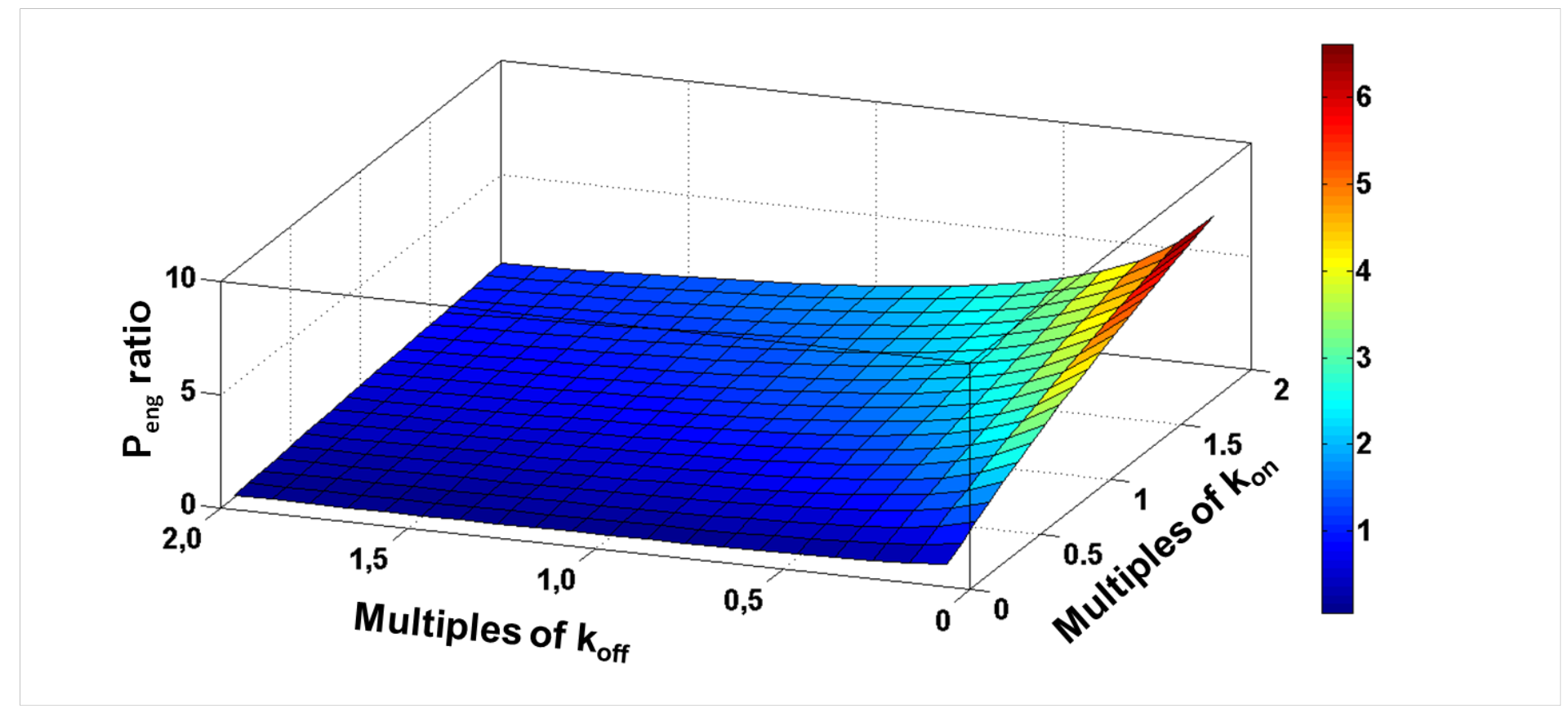
*P*_eng_ fold change as a function of varying k_on_ and k_off_ from one-tenth to two-fold of their original values (i.e., k_on_=0.0216 s^−1^ and k_off_=0.00159 s^−1^).

**Figure S7.**
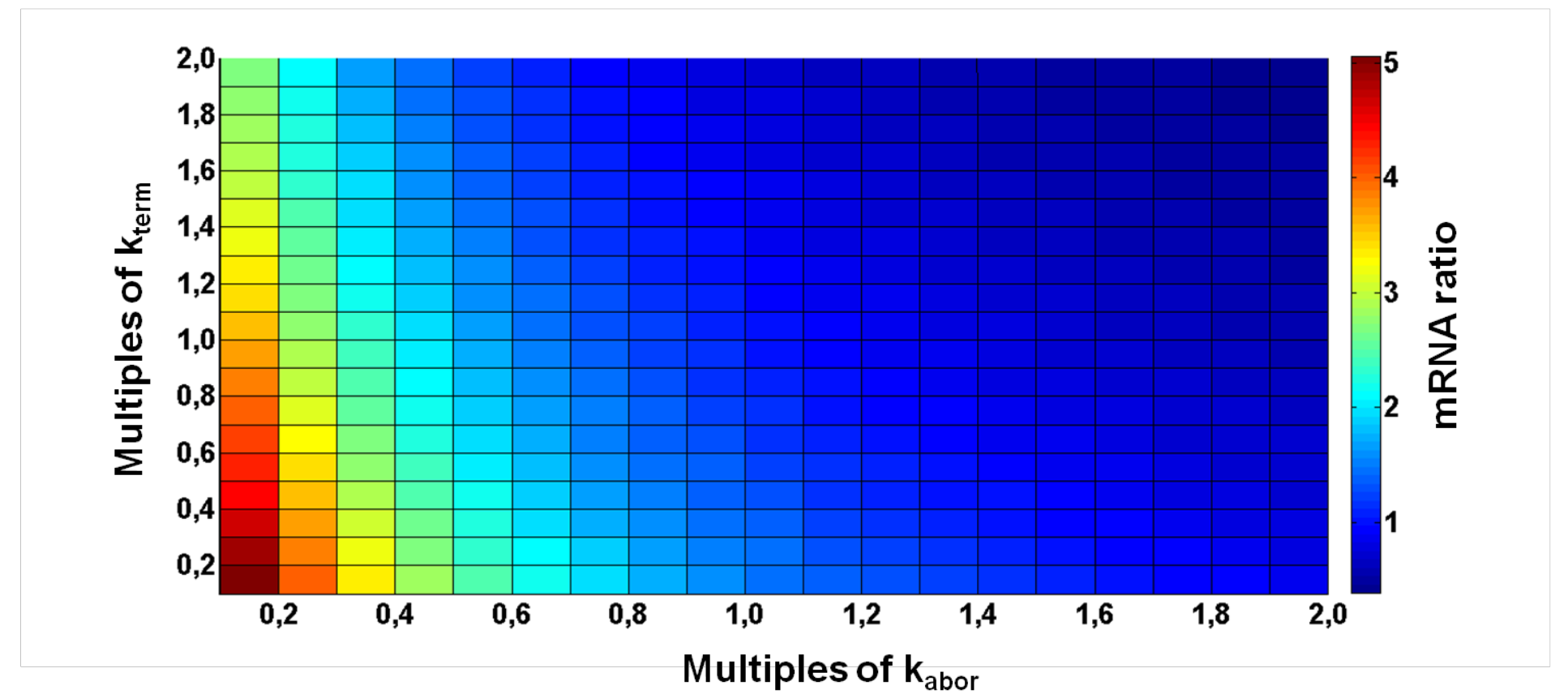
*mRNA* fold change as a function of varying k_abor_ and k_term_ from one-tenth to two-fold of their original values (i.e., k_abor_=0.0170 s^−1^ and k_term_=0.0016 s^−1^).

**Figure S8.**
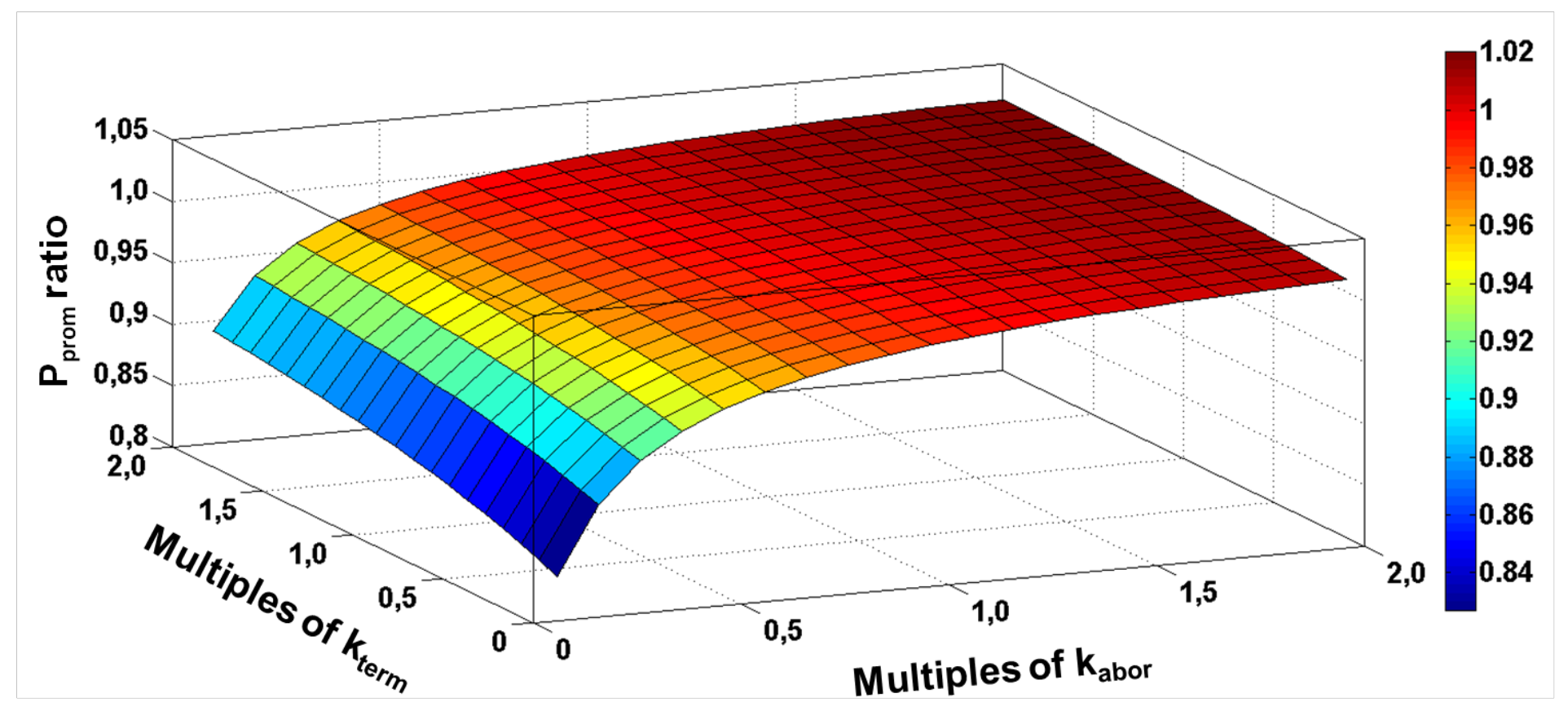
*P*_prom_ fold change as a function of varying k_abor_ and k_term_ from one-tenth to two-fold of their original values (i.e., k_abor_=0.0170 s^−1^ and k_term_=0.0016 s^−1^).

**Figure S9.**
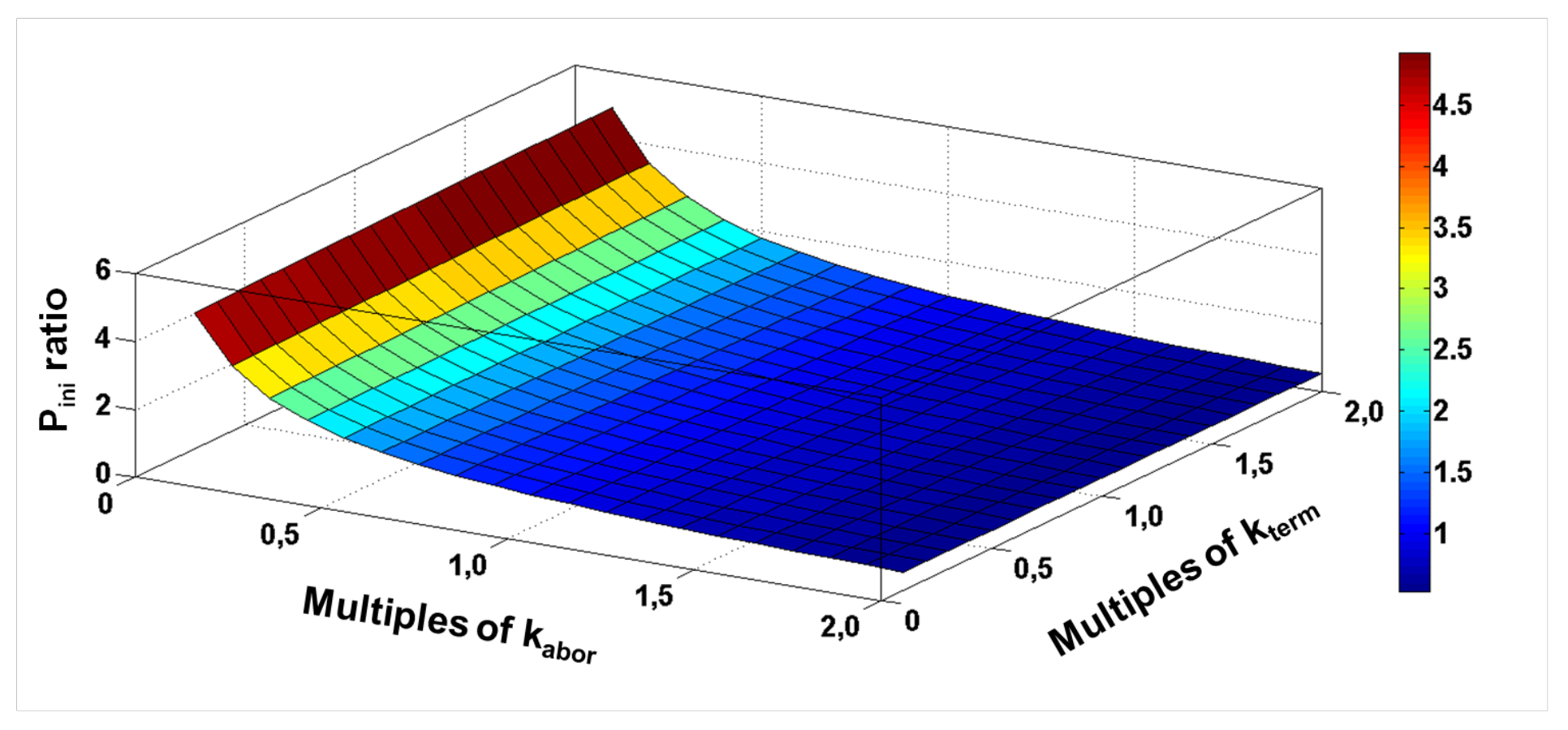
*P*_ini_ fold change as a function of varying k_abor_ and k_term_ from one-tenth to two-fold of their original values (i.e., k_abor_=0.0170 s^−1^ and k_term_=0.0016 s^−1^).

**Figure S10.**
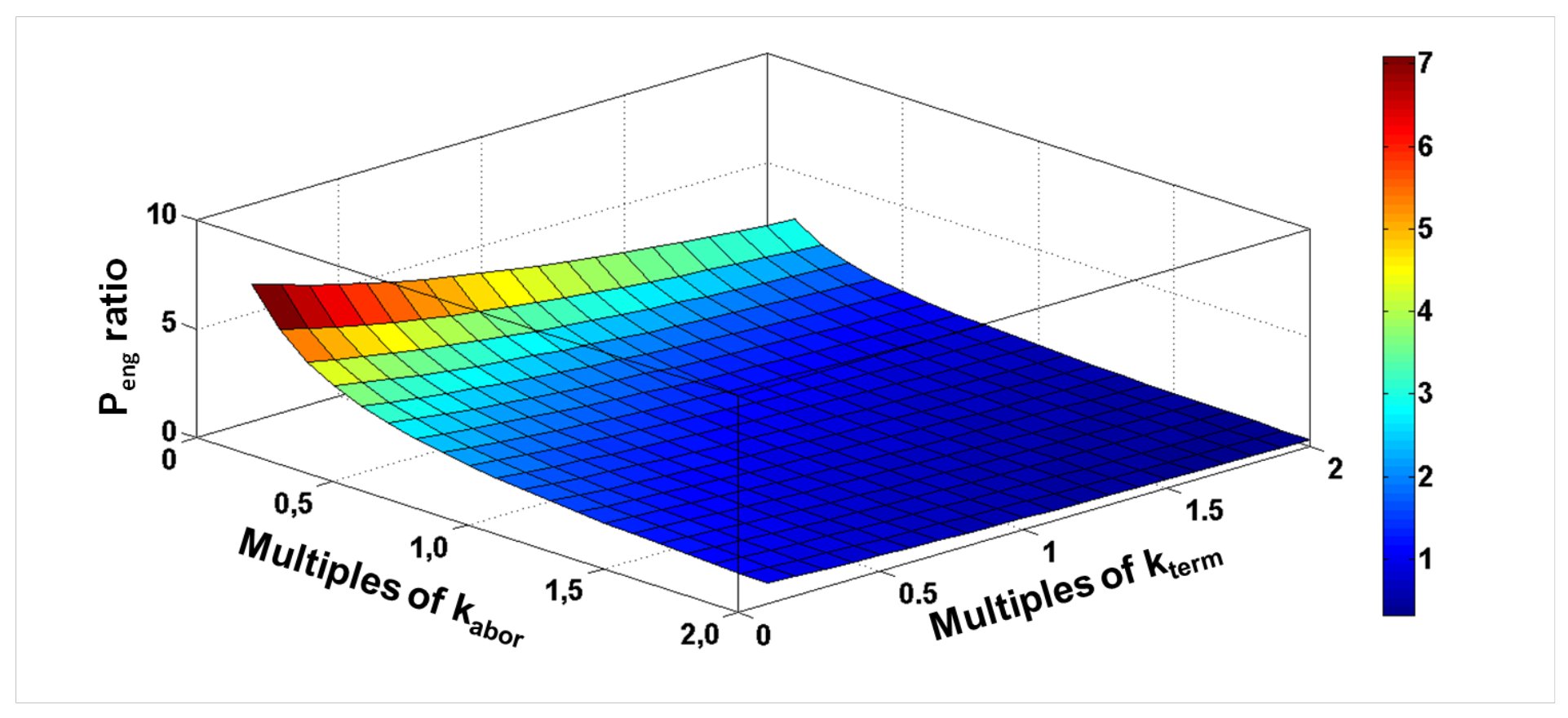
*P*_eng_ fold change as a function of varying k_abor_ and k_term_ from one-tenth to two-fold of their original values (i.e., k_abor_=0.0170 s^−1^ and k_term_=0.0016 s^−1^).

## References

1. Meyer, B.J. X-Chromosome dosage compensation. (2005).

2. Akhtar, A. Dosage compensation: an intertwined world of RNA and chromatin remodelling. Current opinion in genetics & development 13, 161–169 (2003).

3. Conrad, T. & Akhtar, A. Dosage compensation in Drosophila melanogaster: epigenetic fine-tuning of chromosome-wide transcription. Nature reviews. Genetics 13, 123 (2012).

4. Prestel, M., Feller, C., Straub, T., Mitlöhner, H. & Becker, P.B. The activation potential of MOF is constrained for dosage compensation. Molecular cell 38, 815–826 (2010).

5. Hamada, F.N., Park, P.J., Gordadze, P.R. & Kuroda, M.I. Global regulation of X chromosomal genes by the MSL complex in Drosophila melanogaster. Genes & development 19, 2289–2294 (2005).

6. Straub, T., Gilfillan, G.D., Maier, V.K. & Becker, P.B. The Drosophila MSL complex activates the transcription of target genes. Genes & development 19, 2284–2288 (2005).

7. Park, S.-W., Oh, H., Lin, Y.-R. & Park, Y. MSL cis-spreading from roX gene up-regulates the neighboring genes. Biochemical and biophysical research communications 399, 227–231 (2010).

8. Ohno, S., Kaplan, W. & Kinosita, R. Formation of the sex chromatin by a single X-chromosome in liver cells of Rattus norvegicus. Experimental cell research 18, 415–418 (1959).

9. Lyon, M.F. Gene action in the X-chromosome of the mouse (Mus musculus L.). nature 190, 372–373 (1961).

10. Gartler, S.M. & Riggs, A.D. Mammalian X-chromosome inactivation. Annual review of genetics 17, 155–190 (1983).

11. Disteche, C.M. Dosage compensation of the active X chromosome in mammals. Nature genetics 38, 47 (2006).

12. Yildirim, E., Sadreyev, R.I., Pinter, S.F. & Lee, J.T. X-chromosome hyperactivation in mammals via nonlinear relationships between chromatin states and transcription. Nature structural & molecular biology 19, 56–61 (2012).

13. Deng, X. et al. Mammalian X upregulation is associated with enhanced transcription initiation, RNA half-life, and MOF-mediated H4K16 acetylation. Developmental cell 25, 55–68 (2013).

14. Larschan, E. et al. X chromosome dosage compensation via enhanced transcriptional elongation in Drosophila. Nature 471, 115 (2011).

15. Conrad, T., Cavalli, F.M., Vaquerizas, J.M., Luscombe, N.M. & Akhtar, A. Drosophila dosage compensation involves enhanced Pol II recruitment to male X-linked promoters. Science 337, 742–746 (2012).

16. Ferrari, F. et al. Comment on “Drosophila dosage compensation involves enhanced Pol II recruitment to male X-linked promoters”. Science (New York, N.Y.) 340, 273–273 (2013).

17. Vaquerizas, J.M., Cavalli, F.M., Conrad, T., Akhtar, A. & Luscombe, N.M. Response to comments on “Drosophila dosage compensation involves enhanced Pol II recruitment to male X-Linked promoters”. Science 340, 273–273 (2013).

18. Ferrari, F. et al. “Jump start and gain” model for dosage compensation in Drosophila based on direct sequencing of nascent transcripts. Cell reports 5, 629–636 (2013).

19. Darzacq, X. et al. In vivo dynamics of RNA polymerase II transcription. Nature structural & molecular biology 14, 796–806 (2007).

20. Ehrensberger, A.H., Kelly, G.P. & Svejstrup, J.Q. Mechanistic interpretation of promoter-proximal peaks and RNAPII density maps. Cell 154, 713–715 (2013).

21. Henriques, T. et al. Stable pausing by RNA polymerase II provides an opportunity to target and integrate regulatory signals. Molecular cell 52, 517–528 (2013).

22. Chlamydas, S. et al. Functional interplay between MSL1 and CDK7 controls RNA polymerase II Ser5 phosphorylation. Nature 201, 6 (2016).

23. Kim, J.K. & Marioni, J.C. Inferring the kinetics of stochastic gene expression from single-cell RNA-sequencing data. Genome biology 14, R7 (2013).

## References

1 Ferguson, Heather A., Jennifer F. Kugel, and James A. Goodrich. “Kinetic and mechanistic analysis of the RNA polymerase II transcription reaction at the human interleukin-2 promoter”. Journal of molecular biology 314.5 (2001): 993–1006.

2 Darzacq, Xavier, et al. “In vivo dynamics of RNA polymerase II transcription.” Nature structural & molecular biology 14.9 (2007): 796–806.

